# A framework for modelling desert locust population dynamics and large-scale dispersal

**DOI:** 10.1101/2023.07.11.548524

**Authors:** Renata Retkute, William Thurston, Keith Cressman, Christopher A. Gilligan

## Abstract

As climate change leads to new areas being under threat from emerging pests, there is increasing demand for approaches combining mechanistic models and geospatial information. Forecasting based on modelling can inform surveillance, early warning and management. This paper introduces a novel framework for modelling desert locust population dynamics and migration using knowledge of pest biology, climate and remote sensing data, wind trajectories and desert locust survey data from Locust Hub (FAO). Furthermore, we propose an algorithm for forecasting short-term and long-term desert locust migration based on limited reporting data.

## 1. Introduction

Desert locust (*Schistocerca gregaria*) is one of the most dangerous migratory pests for agricultural and rangeland production (Lecoq, 2003), making control a priority for food security in many regions (Retkute et al., 2021). The period 2019–2021 was marked by one of the largest desert locust upsurges over two decades, which saw desert locust (DL) invasion extending from Kenya to India (Salih et al., 2020; Lecoq and Cease, 2022). Mathematical models have previously been developed and used to assess particular aspects of locust dynamics. These include: collective motion of gregarious locusts (Yates et al., 2009; Bazazi et al., 2010; Ariel and Ayali, 2015; Bernoff et al., 2020), longitudinal flight dynamics (Taylor, 1979; Taylor and Zbikowski, 2005), foraging behaviour Georgiou et al. (2021, 2022), phase transitions between solitarious and gregarious forms (Holt and Cheke, 1996; Topaz et al., 2012; Hassanali et al., 2005; Akimenko and Piou, 2018), forecasting gregarization areas (Veran et al., 2015), population dynamics (Cheke, 1978; Farrow and Longstaff, 1986; Ibrahim, 2008), and influence on crop production (Mamo and Bedane, 2021a,b). Recently, data analytics methods, such as regression and machine learning, have been applied to predict breeding areas of DL (Piou et al., 2013; Gómez et al., 2018; Kimathi et al., 2020; Ellenburg et al., 2021; Klein et al., 2022), potential habitat areas of DL Meynard et al. (2017); Guan et al. (2021); Saha et al. (2021); Youngblood et al. (2022), presence of DL (Tratalos et al., 2010; Piou et al., 2019; Samil et al., 2020; Tabar et al., 2021; Gómez et al., 2021a; Shao et al., 2021; Yusuf et al., 2021; Sun et al., 2022; Rhodes and Sagan, 2022; Cornejo-Bueno et al., 2022) and the incidence of locust outbreaks ((Lawton et al., 2022)). They have also been applied to detect DL impact on vegetation senescence (Adams et al., 2021), and cropland damage (Alemu and Neigh, 2022). Major challenges remain in spatiotemporal forecasting of DL, in particular to predict the long–range movements of swarms. The dispersal ability of swarms is a key element in assessing the risks to crop and pasture land and in optimising the management of DL. (Zhang et al., 2019; Piou and Marescot, 2023).

The current practice for predicting swarm movement relies heavily on expert opinion to integrate data from surveillance, past knowledge of DL behaviour, current and predicted wind movements. Since its inception by FAO approximately 50 years ago, the desert locust early warning system has been successful in reducing the severity and impacts of DL upsurges during that period Cressman (1996); Healey et al. (1996); Cressman (2016). New technologies including computational modelling, remote sensing and crowd-sourcing are being progressively incorporated into the FAO early warning system for DL. During the recent 2019-21 DL upsurge, near real–time outputs from weather-driven atmospheric transport models, adapted to model DL swarm dispersal, were used to improve predictions of areas at risk. These innovations included twice-weekly forecasts of likely swarm movements using NAME (the Numerical Atmospheric-dispersion Modelling Environment, Jones et al. (2007)) provided by the UK Met Office (Meyer et al., 2023) and a web–based app for forward or backward simulation of swarm movement based on air trajectories from the Hybrid Single-Particle Lagrangian Integrated Trajectory (HYSPLIT) model, provided by the US National Oceanic and Atmospheric Administration (NOAA) (NOAA)). The availability of atmospheric transport models for wind-assisted dispersal of DL provides a valuable tool for analysis (using historic weather data) and prediction (using forecast weather and surveillance data) of long-distance swarm movements. Atmospheric transport models have previously been used to analyse the migratory trajectories of insects, including the annual migration of painted lady butterflies (Hu et al., 2021), seasonal migration of fall armyworm moths (Westbrook et al., 2015) and midge flight activity (Burgin et al., 2012). The NAME model also underpins a successful, near real-time, weather-driven, early warning system for long-range dispersal of wheat rust spores (Meyer et al., 2017) initially deployed in Ethiopia (Allen-Sader et al., 2019) and now extended to South Asia. In the current paper, we position the central role of swarm dispersal within a modelling framework that integrates breeding and development from egg, hopper-to-swarm emergence, migration and feeding. The framework builds on epidemiological modelling concepts (Gilligan, 2008) and an atmospheric transport model for swarm movement. The framework takes account of surveillance data for input and validation, as well as dynamically changing conditions that affect the availability of vegetation for locust feeding. It is designed for analysis and prediction at country-wide and regional scales, with a spatial resolution of 1km *×* 1km .

A principal aim of the framework is to provide a practical starting point for use in the next upsurge. Specifically we address the following topics:

- construction of a modelling framework that integrates DL breeding, development through egg, hopper and adult stages, feeding and swarm migration;
- characterisation of breeding sites using a combination of site-specific static and dynamic variables to predict egg-laying, hopper and adult development leading to DL swarming;
- incorporation of weather-driven models for wind trajectories to predict daily pathways of swarm migration;
- use of remote-sensed data to predict the duration of swarm feeding at a single location given the state and availability of vegetation for feeding;
- use of the modelling framework to analyse and predict: (i) long–distance, seasonal movement of swarms, e.g. to reach Kenya from breeding sites in Somalia; (ii) short–term daily and weekly movements of swarms.

We also address the question of how to assess flight direction and likely origin and landing sites for swarms observed in flight.

## 2. Methods

### 2.1. Data on DL surveys, environment and remote sensing

Historical data on desert locust presence and absence were obtained from survey reports and archives at the FAO Locust Hub website (FAO). The data consist of reported locations for adults (solitarious), hoppers (solitarious), hopper bands (gregarious) and adult swarms (gregarious). We extracted the records dated for the period 1 January 2019 and 31 December 2021 that covers the recent DL upsurge in East Africa.

We extracted data for land cover classification from the Copernicus global map of land cover at 100 m resolution (CLC100) (Buchhorn et al., 2020). Land cover maps represent spatial information on different types of the Earth’s surface, e.g. urban areas, sparse vegetation, cultivated vegetation. For NDVI values, we used 8-day averaged moderate-resolution imaging spectroradiometer (MODIS) data (Terra Surface Reflectance 8-Day Global 250m, MOD09Q1 v006) (Vermote, 2015). Other data sets we used were: elevation (the Shuttle Radar Topography Mission (SRTM) digital elevation dataset) (Jarvis et al., 2008 (Accessed: 1 June 2022); sand content in the soil at 5cm depth (weight percentage of the sand particles (0.05–2 mm) from ISRIC soil-grids dataset) (Hengl et al., 2017; ISRIC, 2017 (Accessed: 1 June 2022); and clay content in the soil at 5cm depth (weight percentage of the clay particles (*<*0.0002 mm) from ISRIC soilgrids dataset) (Hengl et al., 2017; ISRIC, 2017 (Accessed: 1 June 2022). More details on input data (type, scale, source and provider) are given in Supplementary Table 1.

### 2.2. NAME model forecasting of trajectories and climate conditions

The Lagrangian atmospheric dispersion model NAME (Jones et al., 2007) was used to calculate stochastic trajectories of potential wind-borne DL swarm migrations. Originally developed by the UK Met Office to model the release, transport, dispersion and removal of hazardous material in the atmosphere (Jones et al., 2007), NAME has subsequently been adapted to model long-distance wind-borne transport of insect vectors of viral disease (Burgin et al., 2012) and fungal spores (Meyer et al., 2017). We calculated stochastic ensembles of wind-borne DL swarm trajectories, originating from 3860 different source locations spaced on a regular 20km *×* 20km grid across Ethiopia, Somalia, Eritrea and Kenya. For every day between 1st September 2019 and 31st December 2021, 1000 individual trajectories were calculated from each of the 3860 starting locations, each commencing 2h after local sunrise and terminating 1h before local sunset to reflect the diurnal flight behaviour of DL swarms (FAO, 2021 (Accessed: 1 June 2022). This resulted in the generation of nearly 3.3 billion trajectories over the study period, each providing highly-resolved data on swarm position (longitude, latitude and altitude) at a temporal resolution of 0.5h. These NAME simulations were driven using historic analysis meteorology from the global configuration of the Unifed Model (UM), the Met Office’s operational Numerical Weather Prediction model (Walters et al., 2019). The UM also provided the surface precipitation rate, near-surface air temperature and 0–10 cm soil moisture at 3 hourly time interval on its native horizontal grid of approximately 10 km by 10 km. Grid is shown on lower left side of Fig. 1. More details on input data (type, scale, source and provider) are given in Supplementary Table 1.

**Figure 1:**
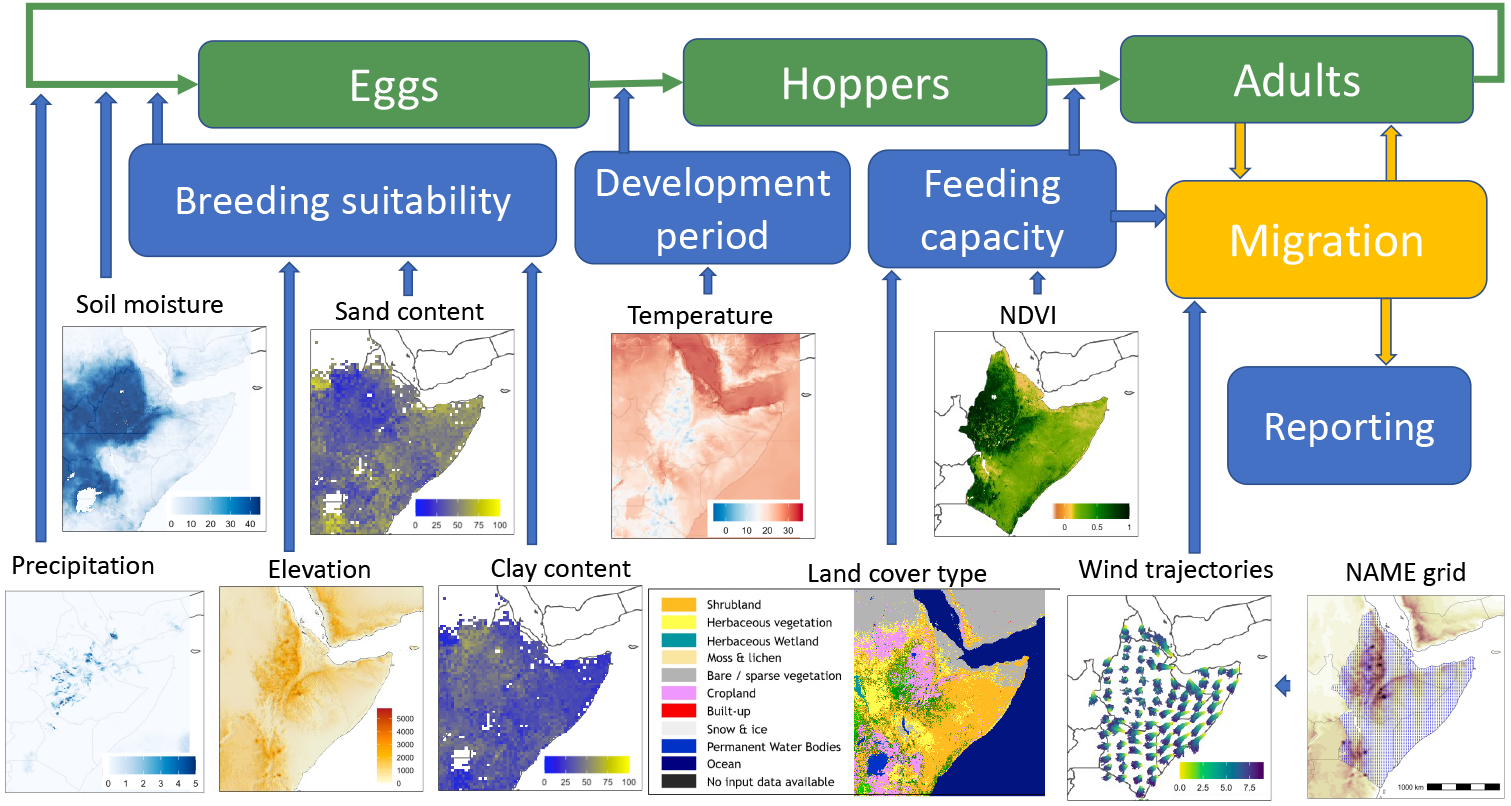
Conceptual diagram of the spatially-explicit model that integrates the life cycle of desert locust, climate variables, remote sensing data, and wind–assisted migration. The model operates on a gridded landscape at 1km *×* 1km resolution. The model is restricted to the dynamics of gregarious locusts.

### 2.3. Mapping breeding suitability

We applied a machine learning approach to combine environmental data with reported locust occurrence and absence in order to predict areas suitable for breeding. We aggregated hopper and band occurrence records (species presence) with DL absence records which had no DL presence within a 10km radius (species absence) and used these data as a response variable. Similar concepts of predicting breeding regions by correlating a set of environmental conditions to presence and absence of DL have been applied previously using a range of factors, for example, temperature, rainfall, sand and moisture contents (Kimathi et al., 2020); soil moisture and soil texture (Ellenburg et al., 2021), or using soil moisture time series only (Gómez et al., 2021b). We restricted our analysis to using three static variables: elevation; sand content in the soil at 5cm depth; and clay content in the soil at 5cm depth. Breeding suitability mapping calculations were performed on the Google Earth Engine platform (Gorelick et al., 2017). We used a Random Forest (Ho, 1995) classifier with the following options: the number of decision trees was set to 100, the number of variables per split was set to 2, the minimun leaf population (i.e. the end node) was set to 2, and the initial seed was set to 0. The im-portance of covariates was obtained by normalising the classifier importance values. The breeding suitability values were adjusted to a 0-100 scale.

We compared our predicted breeding suitability for Kenya with predicted breeding probabilities by Kimathi *et al*. Kimathi et al. (2020) in order to assess the broad consistencies of the approaches. Kimathi *et al*. used datasets at a spatial resolution of 1 km, with the exception of soil moisture, which had resolution downscaled from approximately 55km to 1km. Figure 5A from (Kimathi et al., 2020) was converted into geotiff and values extracted using the R package **raster** (Hijmans, 2022). For ease of comparison, we followed the same breeding suitability classification in five classes as Kimathi et al. (2020): *<* 20, 20-40, 40-60, 60-80 and *>* 80.

### 2.4. Egg laying

We start our simulations from the transition from adults to eggs (breeding). For a sampled location we determine if the location is suitable for breeding by drawing a number from a random uniform distribution and testing if this number is smaller than the breeding suitability value for the given location. Ellenburg *et al*. Ellenburg et al. (2021) showed that soil texture combined with the modelled weekly average volumetric soil moisture from the top layer (0–10 cm) were good predictors of the conditions suitable for locust breeding sites. We combined our predicted suitability estimates with a requirement for soil moisture to be above 12 mm at any time 24-48 hours prior egg laying (Ellenburg et al., 2021; Cressman, 2013).

### 2.5. Development from eggs to hoppers

We model two main factors that influence the rate and success of egg development: temperature and soil moisture Cressman (2013). Egg incubation period can range between 10-65 days FAO (2021 (Accessed: 1 June 2022) and is temperature dependent Cressman (2013). The percentage daily development can be expressed as a function of temperature (in Celsius) Symmons and Cressman (2001):

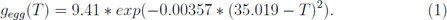

The weekly average of soil moisture required for DL eggs to remain viable was estimated to be between 0.12 and 0.27 *cm*^3^*/cm*^3^ Ellenburg et al. (2021). If soil moisture were outside this range at any week during egg development, we assumed that the eggs had not hatched into hoppers.

### 2.6. Development from hoppers to adults

We assumed that once hoppers hatch from eggs, their movement is limited and contained within the 10km x 10km grid cell they inhabit, which is consistent with reported movements of hopper bands (FAO, 2021 (Accessed: 1 June 2022). Hopper development takes 36 days on average (range 24-95 days) (Symmons and Cressman, 2001). We sampled the length of the development period from a distribution of 24 days + lognormal with mean 2.1 and standard deviation 1, which was truncated between 24 days to 95 days. This gives an asymmetric distribution with a mean of around 36 days and a long tail with a small probability that development would exceed 50 days. The success of the development from hoppers to adults depends on habitat conditions (FAO, 2021 (Accessed: 1 June 2022). Hoppers prefer newly developed vegetation (Pekel et al., 2011). The normalized difference vegetation index (NDVI) is a remote-sensing based index that has high correlation with many photosynthetic characteristics of plants and is used to identify vegetation state (Pettorelli, 2013). As remote sensing data are usually noisy due to cloud cover and other factors (Cai et al., 2017), we applied a Whit-taker smoother for NDVI time series analysis (Frasso and Eilers, 2014; Eilers et al., 2017). Environmental conditions for hopper development were deemed to be suitable in our model if either of two conditions were satisfied during the hopper development period: (i) smoothed NDVI was increasing; or (ii) smoothed NDVI was above a threshold, which we set to be 0.09 Despland et al. (2004).

### 2.7. Calculation of NDVI trend

The NDVI trend was estimated in two steps. First, we used the **R** package *segmented* Muggeo (2017) to calculate break-points of the smoothed NDVI curve. Then we combined segments that had the same direction of change, i.e. increasing NDVI value or decreasing NDVI value. The last segment before swarm landing date then was used to define the NDVI trend (increasing, constant or decreasing); the length of the segment window in days; NDVI value at the start of the segment (when the trend started); and the NDVI value at the time of swarm landing.

### 2.8. Characterising the ensemble of dispersal trajectories by fitting the von Mises distribution

For a given source occupied by a migrating DL swarm on a given day, we characterised the ensemble of wind-assisted dispersal DL trajectories from the site using the von Mises distribution. We first calculated the angle for each trajectory between the East direction and a vector coordinate between the start and end location. The von Mises distribution function was fitted using a modified Adaptive Importance Sampling algorithm (Retkute et al., 2021). We chose uniform priors: *µ ∼ U* [0, 360] (the directional mean), and *κ ∼ U* [0, 700] (concentration of the distribution). The corresponding importance weights are calculated as:

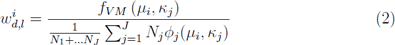

where *f_V M_* (*µ, κ*) is the probability density of the von Mises distribution, *d* is day, *l* is location, *i* is the sampled parameter set, and *ϕ_j_*(*µ, κ*) is a proposal distribution at iteration *j*. The proposal distribution is equal to the prior at the first iteration, and composed as a mixture of Student’s t-distributions for subsequent iterations (Retkute et al., 2021). We ran 10 iterations with 1000 parameter sets sampled each iteration. For analysis, we used a maximum a posteriori (MAP) estimate for each day and location.

### 2.9. Simulation of wind-borne movements of individuals swarms

When simulating dispersal of a swarm, we sampled a single NAME wind trajectory. If coordinates of the recent landing site were not on the grid, we searched for the closest grid coordinates and then translated the corresponding wind trajectory to start at the required location. We assumed that the propensity of a swarm to start flying in upcoming days was dependent on the ground conditions. Wind trajectories leading to the ocean were stopped at the closest grid cell on land. All trajectories that went outside the region shown in Figure 1 were terminated.

## 3. Results

### 3.1. The modelling framework

We consider a domain extending across five sub-Saharan countries (Kenya, Ethiopia, Somalia, Eritrea and Djibouti) that were affected by the 2019-21 DL upsurge. We formulated a spatially explicit and stochastic compartmental framework that follows gregarious locust populations through egg, hopper and adult stages in time and space as they develop, mature, and disperse in search of areas suitable for feeding and breeding within the domain shown in Figure (1). We model the DL swarm as a coherent unit of gregarious locusts at a similar developmental stage and at close proximity (rather than individual locusts). In accordance with our study area, our simulations were restricted to longitude 30 to 55 degrees and latitude between −5 to 20 degrees. We assumed that DL populations can be in one of three non-overlapping life-stages, also called compartments: eggs, hoppers (nymphs), and adults. Each compartment has different ecological requirements and responds differently to environmental conditions, such as climate, soil and vegetation (FAO, 2021 (Accessed: 1 June 2022). The model operates on a gridded landscape at a resolution of 1km *×* 1km over one day time steps and represents three main processes: (i) the biology and behaviour of DL; (ii) meteorological and environmental conditions; and (iii) wind–assisted migration of DL swarms. We restricted the model to dynamics of gregarious locusts, as these pose the largest threat to livelihoods (FAO, 2021 (Accessed: 1 June 2022). As the model focuses on gregarious swarm behaviour and movement, we have not accounted for transitions between solitarious and gregarious phases observed in DL.

The main assumption for DL that we make in the model is that long-distance swarm flights follow wind trajectories (Rainey, 1951). We also assume that swarms take-off under favourable conditions 2h after local sunrise and follow wind trajectories until landing 1h before sunset (FAO, 2021 (Accessed: 1 June 2022). The duration that swarms remain at a landing site is determined by the availability of a suitable food source and we model this duration as a function of land cover type and NDVI.

### 3.2. Areas suitable for breeding

Desert locust outbreaks start with locusts taking advantage of environmental conditions suitable for breeding. Identifying breading areas is impor-tant for both monitoring situations for preventive management and understanding potential DL migratory patterns. Previously, a number of studies derived suitability maps based on a combination of static and dynamic co-variates (Gómez et al., 2018; Kimathi et al., 2020; Ellenburg et al., 2021; Klein et al., 2022). Climate data used were either averaged monthly values based on long term data (Kimathi et al., 2020; Klein et al., 2022), or statistical properties (average, minimum, or maximum) of 6-12 days distribution of daily values (Gómez et al., 2018; Ellenburg et al., 2021). We wanted to take advantage of the high temporal resolution data on precipitation and soil moisture provided by the UM. We separated the breeding process into two parts: derivation of a suitability map, which depends on static covariates (elevation, sand and clay content in soil); and allowance for recent (up to 24 hours) precipitation events.

A suitability map for desert locust breeding sites is shown in Figure 2 (A). Using a normalised score of 0-100 our results indicate that 43% of 1km x 1 km grid cells had breeding suitability higher than 60, and 13% probability higher than 80. The fitted model showed that all three covariates had similar importance: with elevation accounting for 32% of the variation, sand content 34%, and clay content 34%.

**Figure 2:**
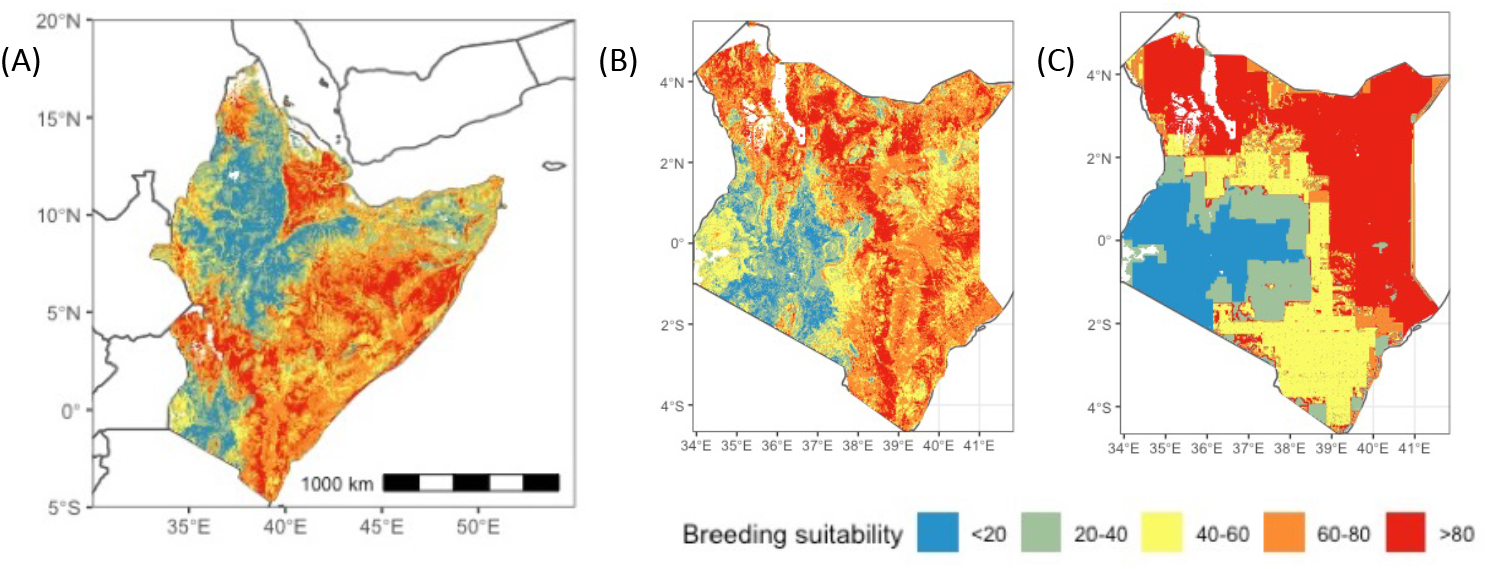
Predicted breeding suitability map : (A) for East Africa -(B) Kenya this study; (C) Kenya, Kimathi *et al*. Kimathi et al. (2020).

We compared our predicted breeding suitability with predictions in (Ki-mathi et al., 2020) (Figure 2 (B)-(C)). We found broad correspondence at high and very high suitability breeding regions between the two models (Figure 2 (B)-(C)), albeit the higher resolution of the model introduced here indicates greater spatial heterogeneity in the suitability scores.

### 3.3. Feeding capacity for swarms

An important aspect of our modelling framework was to assess the effects of vegetation on locust dynamics and on the speed of advance of swarms. After analysing the relationship between land cover types and NDVI, we concluded that three factors determine feeding capacity: land cover type, NDVI trend (increasing, decreasing or constant) and vegetation state (current value of NDVI). We summarised land cover types into four groups: (i) bare/sparse vegetation; (ii) shrubs, herbaceous vegetation and herbaceous wetland; (iii) cultivated and managed vegetation/agriculture (cropland); and (iv) forest. The surface phenology of each land cover group was inferred from the temporal profiles of NDVI (Defries and Townshend, 1994). Depending on the value of the NDVI, the vegetation state on the ground was classified into six classes (Zaitunah et al., 2018): no vegetation (NDVI¡0), lowest density (NDVI*∈* [0, 0.15)), lower density (NDVI *∈* [0.15, 0.3)), dense vegeta-tion (NDVI*∈* [0.3, 0.45)), higher density (NDVI*∈* [0.45, 0.6), highest density (NDVI*∈* [0.6, 1]).

We divided the length of stay of DL swarms into three classes: short stay (1-2 days), medium stay (2-4 days), and long stay (4-7 days) FAO and WMO (2016). Figure 3 (A) shows how we assign the class of stay for swarms depending on environmental conditions at a given location. The number of days a swarm stays and feeds was sampled from the range corresponding to the class to which the location was assigned. For NDVI values less than zero, swarms would stay only over night and start flying in the morning.

**Figure 3:**
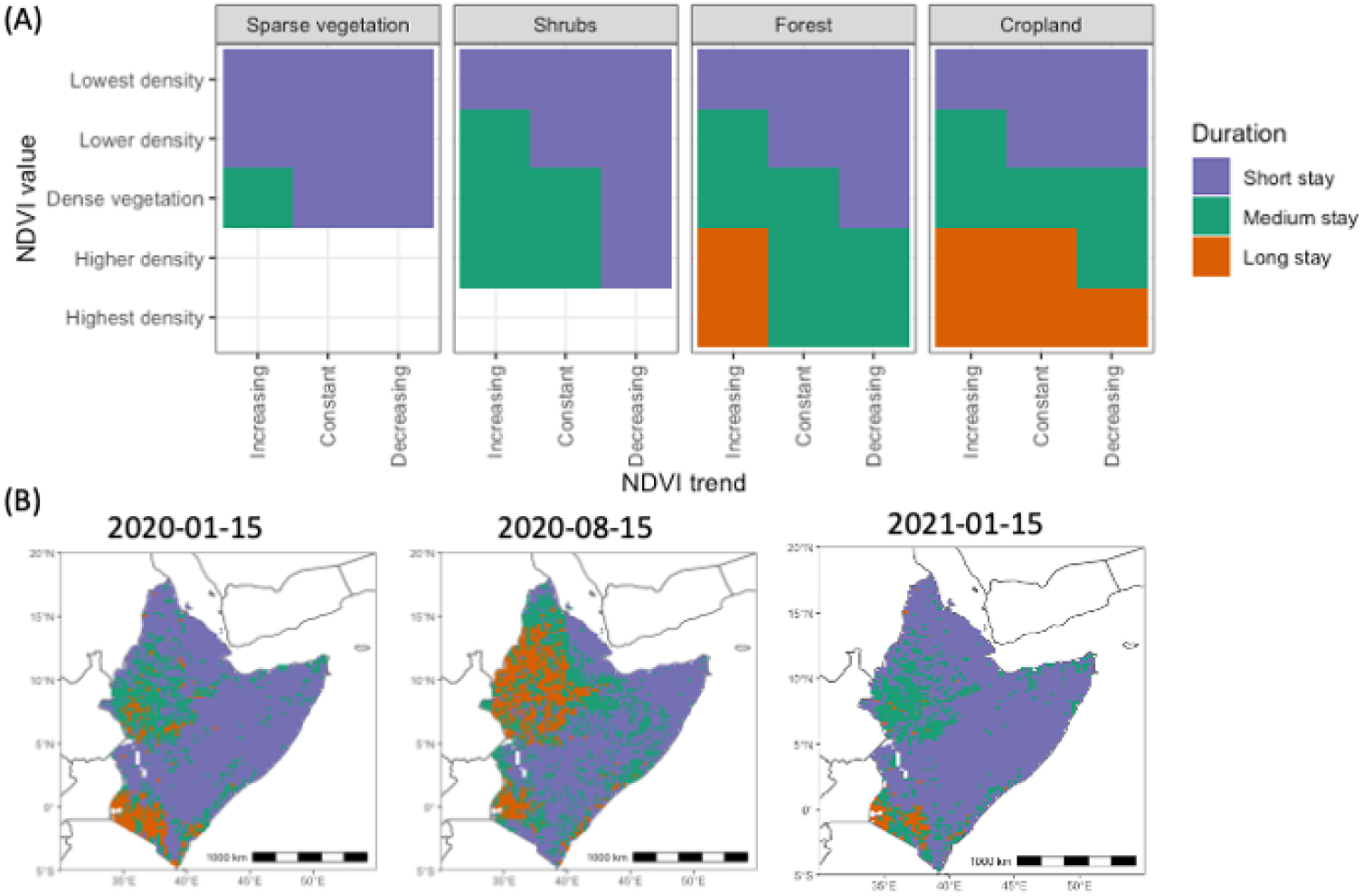
(A) Length of stay for swarms depends on land cover type, NDVI trend (increasing, decreasing or constant) and vegetation state (value of NDVI). Here short stay is 1-2 days, medium stay is 2-4 days, and long stay is 4-7 days. (B) Spatial distribution of stay length. Length of stay was classified as short stay (1-2 days), medium stay (2-4 days), or long stay (4-7 days).

Examples of the spatial distribution of potential length of stay are shown in Figure 3(B) for three dates: 15th of January 2020, 15th August 2020, and 15th of January 2021 for the target domain in East Africa. Most areas were classified as short stay duration on all three dates. Long stay areas on 15th of August 2020 correspond to mid-season of major crop types in Ethiopia (Qu and Hao, 2018).

### 3.4. Variation of wind patterns

The migration pathways achieved by flying insects are highly dependent on the seasonal variation of wind patterns (Cheke and Tratalos, 2007; Chapman et al., 2010; Dingle, 2014). There is stochastic variation in the direction of individual realisations starting from the same location, due to the turbulent nature of the atmosphere as represented in NAME. An example of wind trajectories on the 15th of January 2021 is shown in Figure 4 (A), and a close-up at a particular location in Figure 4 (B). Seasonal variation in wind direction at the selected location can be clearly seen when the distribution of wind direction is plotted as a function of date (Figure 4(D)). For example, south-westerlies dominated from June to October in 2020, switching to north-easterlies from October 2020 to March 2021, consistent with the seasonal reversal of the Asian monsoon winds (Findlater, 1971).

**Figure 4:**
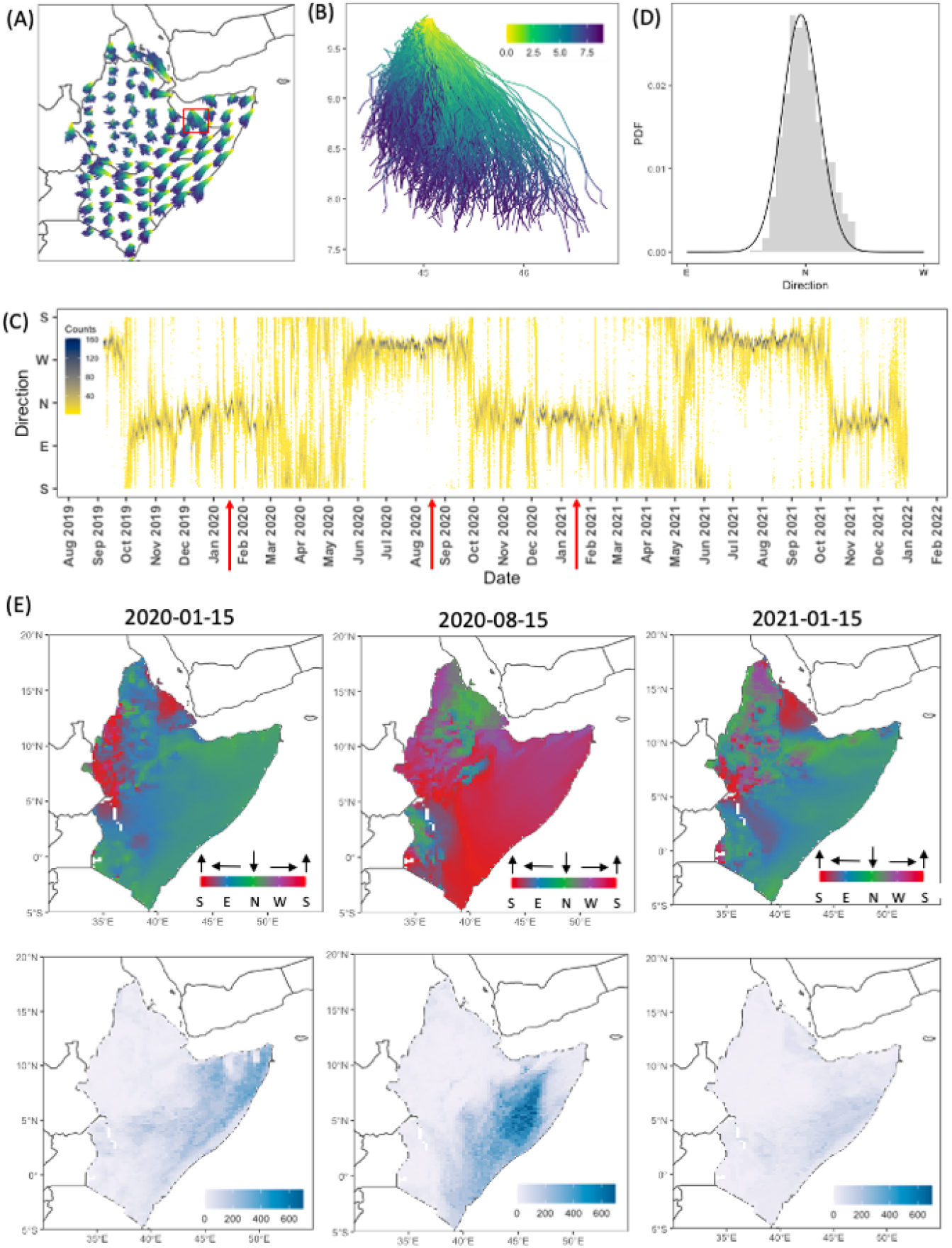
(A) Wind trajectories from a subset of the 3860 starting points on the 15th of January 2021. (B) Wind trajectories from a starting point enclosed in the red square in (A). The colour shows the difference in hours between points on a trajectory and the initial point. (C) Histogram of directional data from (B) (gray) and fitted von Mises distribution (black curve). (D) Daily distribution of directional data from a starting point enclosed in the red square in (A). Direction was divided into 5 degree bins and the colour the number of trajectories in each bin. (E) MAP estimates of fitted the von Mises distribution: directional mean (top row) and concentration (lower row).

We fitted the von Mises distribution to angles representing the directions of NAME wind trajectories. The von Mises distribution is defined by two parameters: the directional mean and concentration of the distribution. The directional mean indicates seasonal variation of global wind patterns, while the concentration can be used as a measure of stochasticity introduced by NAME trajectories to represent natural variation in turbulent wind flow. Values of concentration close to zero indicate a uniform angular distribution, and high values imply low variance that is consistent with stronger directional flow. An example of the fitted distribution is shown in Figure 4 (C), where the histogram is for directional data from trajectories in Figure 4 (B) and the black line is derived from the fitted von Mises distribution. To illustrate global wind trends, we calculated wind trajectory angles and fitted the von Mises distribution for three dates: 15th of January 2020, 15th of August 2020 and 15th of January 2021. The fitted directional mean for January had similar trends for 2020 and 2021, i.e. most of wind trajectories had southwards orientations, but some differences could be observed for certain areas (Figure 4 (E)). The fitted concentration values showed larger variation on 15th January between the two years, with northerly wind concentrated more in 2020 than in 2021. Meanwhile, wind trajectories were directed towards the north on 15th of August 2020, with a larger range of concentrations in comparison with wind trajectories in January for both 2020 and 2021.

### 3.5. The simulation study

Having constructed a modelling framework that integrates DL breeding, development through life cycle stages, feeding and swarm migration (Figure 1) we then assessed the ability of the model to reproduce long-distance movements of DL swarms observed during the 2019-2020 DL upsurge. For this, we extracted data from the Locust Hub DL surveys for two time periods. The first period (1*^st^* September - 1*^st^* October 2020) corresponded to the egg laying time (FAO, 2020), and the second period (1*^st^* December 2020 - 15*^th^* January 20221) corresponded to the observed mass migration of DL from north Somalia and Ethiopia to Kenya (FAO, 2021). The locations of reported swarm presence for the two time periods are shown in Figure 5 (A). Specifically, we simulated breeding hopper and swarm emergence in Somalia during September 2020 and assessed whether or not simulations of subsequent swarm migration and feeding were consistent with reported observations of swarms arriving in Kenya, between four and five months later.

**Figure 5:**
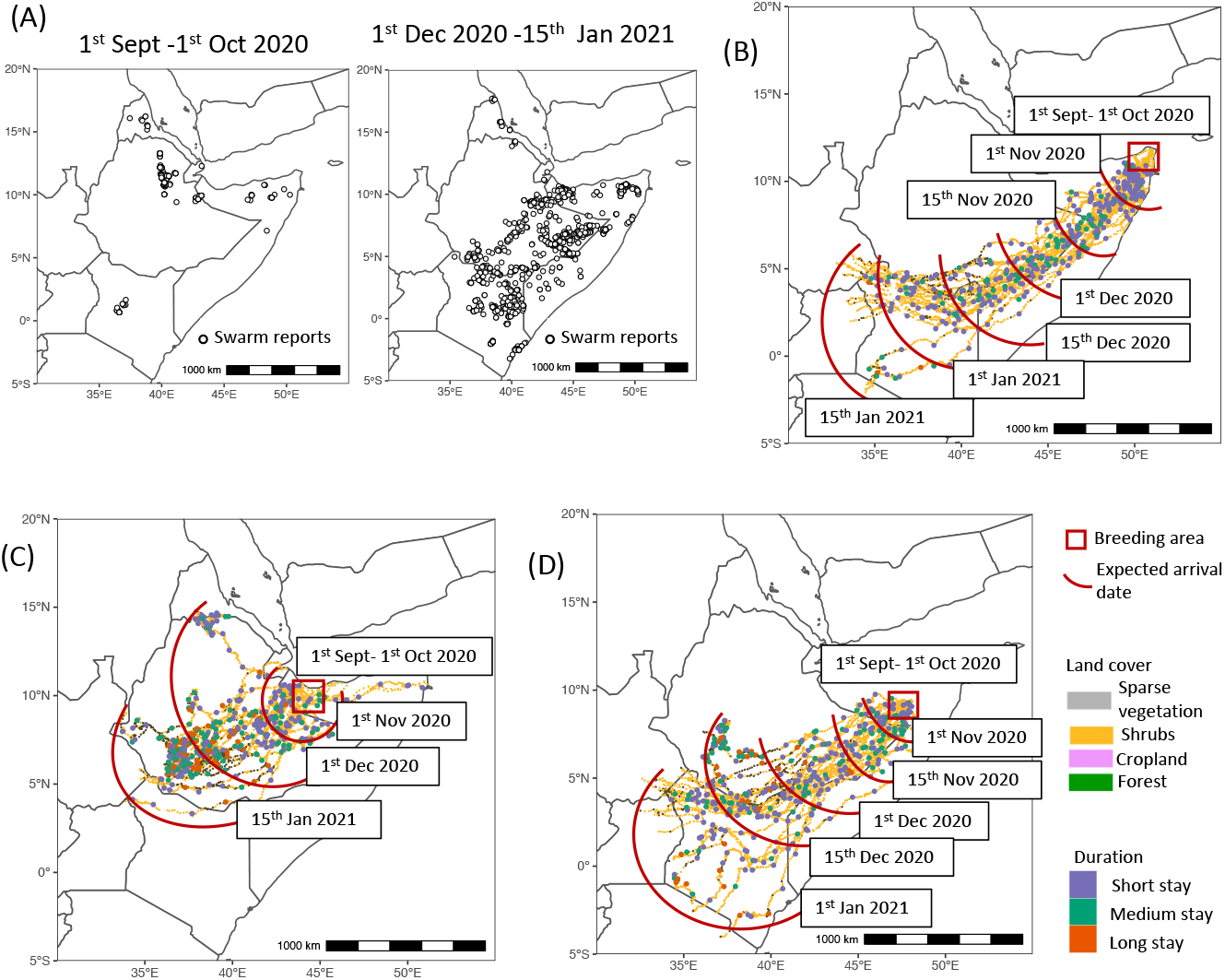
(A) Locations of reported swarms from Locust Hub during two periods. (B)-(D) Simulated breeding, development and migration of DL. Red rectangles indicate areas that were sampled as a breeding ground. Lines show simulated DL migration trajectories, which are coloured according to land cover type at landing sites. Circles indicate landing locations and are coloured according to the duration swarms stayed. Red curves indicate date (on average) when swarms reached a particular area.

We chose a breeding time in September 2020, based on FAO data and reports of the situation in the Horn of Africa FAO (2020). We ran a pilot study to identify representative breeding sites that were consistent with locust surveys. First, we randomly sampled points in northern Somalia as potential breeding locations. Starting from each location we simulated locust breeding, development, feeding and dispersal using historic weather data from 15 September 2020 until 15 January 2021. Simulations that failed to satisfy conditions for development from emergence to adults and simulations for which sampled trajectories terminated in permanent water bodies, were deemed unsuccessful. After visual inspection of generated migration trajec-tories of successful simulations, we selected three representative locations as breeding areas, which are shown as red squares in Figure 5 (B)-(D)). For each selected site, we re-ran simulations, until 25 successful breeding, development and migration histories were obtained.

For all three breeding sites, DL hatched, developed from hoppers into adults and started migration before November 2020. However, migration was faster from breeding sites in Figures 5(A) and (C), than for the site in Figure 5 (B) as the trajectories for swarms from the sites in Figures 5 (A) and (C) encountered more favourable environmental conditions for feeding. Our simulations show that from mid December 2020 to mid January 2021 simulated swarms would have reached the northeast Kenya 5.

### 3.6. Short-term and long-term risk mapping

Here we investigate how the model could be used for short-term (1-2 days) and long-term (up to 7 days) forecasting of swarm migration. To mimic real-life conditions, we assumed that swarms are most likely to be spotted and reported while they are flying between landing sites. Hence, the only data available are the co–ordinate and date of the sighting, with no information about flight direction, which is notoriously difficult for observers on the ground to assess (Draper, 1980), nor is it known where a swarm came from.

To illustrate short-term and long-term forecasting, we chose a single representative simulation of locust development and migration trajectory with locations and timing close to historical data (Fig.5A-B). We assumed that the simulation represented a large swarm. Figure 6(A) shows a breeding site (location 1), migration trajectory (dashed line) and daily landing sites (locations 2-12). Most of the landing sites were classified as shrubs, and the rest as sparse vegetation.

**Figure 6:**
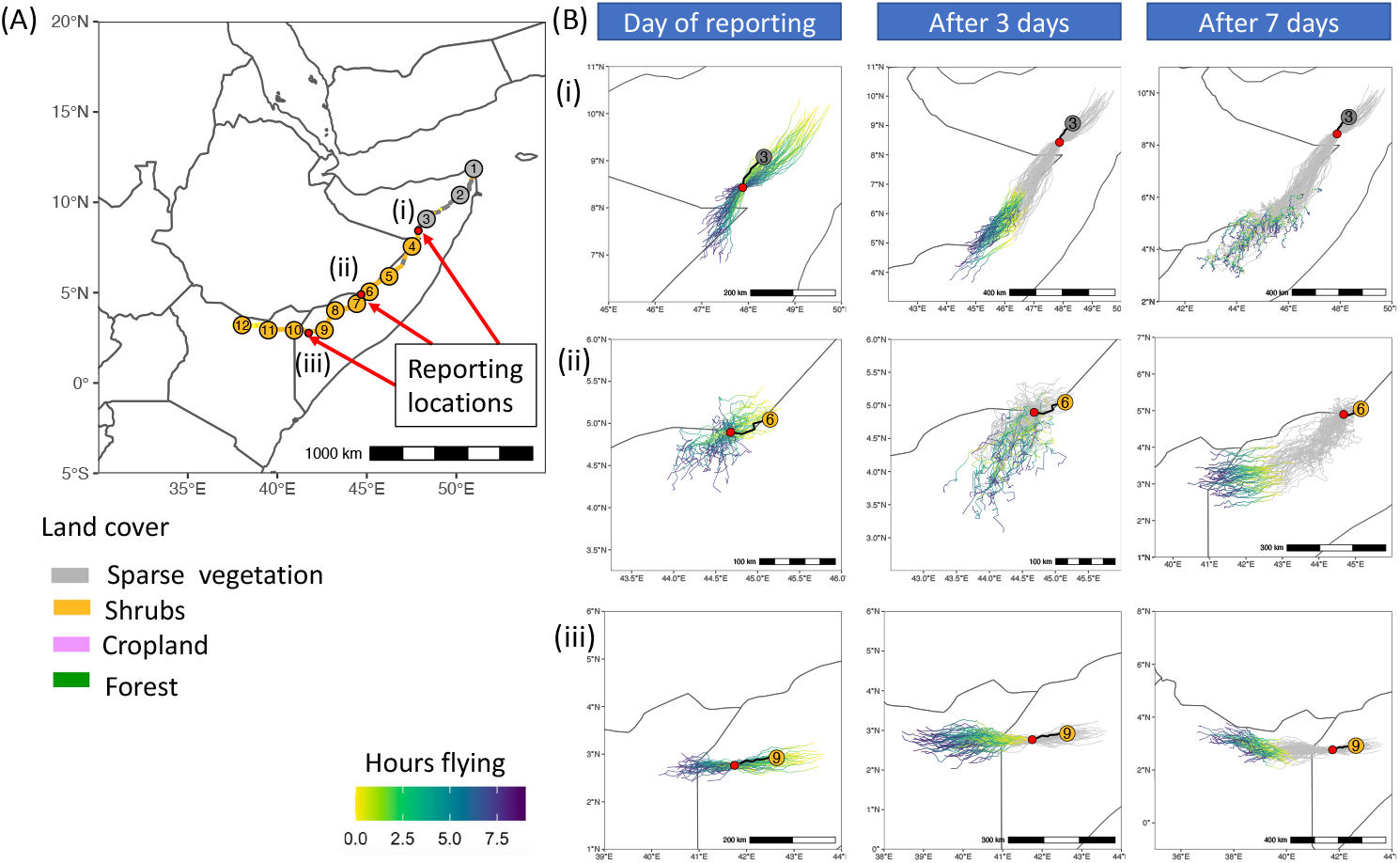
(A) Simulation of locust activity, reporting and forecasting from 15th of September 2020 to 2nd of January 2021: interpretation of a simulated trajectory. Circles show locust landing sites. The breeding and hopper development site is location 1. Circle colours indicate land cover types. Red dots show three reporting locations. (B) Simulation of swarm reporting (red dots) and potential trajectories passing close to the reporting locations. Trajectories on the last flying day are coloured according to hours flying, and trajectories from previous days are shown in gray. Rows correspond to three reporting locations.

We looked at three reporting locations: at the border between Somalia and northern Ethiopia (site *i*, between locations 3 and 4 in Fig.6A), at the border between Somalia and eastern Ethiopia (site *ii*, between locations 6 and 7 in Fig.6A), and at the border between Somalia and Kenya (site *iii*, between locations 9 and 10 in Fig.6A)).

The forecasting process was simulated as follows. First we sampled 10,000 flight start coordinates within a 250km radius around the reporting site. We sampled a trajectory from each starting coordinate for the day of reporting. We retained only those trajectories that passed within 1km or closer to the reporting location. This provided a short-term forecast of potential landing sites (left column in Fig.6 B). The landing points of these trajectories were used to derive a risk map of swarm locations at the end of the reporting day. The model can be used to simulate locust migration and feeding for up to seven days in order to mimic a seven-day (i.e. longer-term) forecast (middle and right columns in Fig.6 B). Here we used historic analysis weather data from the UM for the appropriate locations and times. The models can also be run in near-real time using weather forecast data from the UM.

Our simulations show that for some days there was large heterogeneity in the spatial distribution of predicted landing points, e.g. after 7 days for reporting location (i) (Fig.6) or on the day of reporting and after 3 days for reporting location (ii) (Fig.6). These scenarios correspond to smaller concentrations of wind trajectories (Section 3.4).

## 4. Discussion

We have developed a framework to predict and analyse the population dynamics and large-scale dispersal of desert locust, motivated by the recent upsurge in East Africa. The key features of our model are identification of suitable breeding sites, which is combined with egg and hopper development, and swarm movement, which is influenced by turbulent wind flow. The model also accounts for feeding behaviour of DL, where the duration swarms spend at a landing site is determined by the availability of food at the site.

Previous modelling studies on DL relied predominantly on statistical approaches to analyse time series data and lacked explicit mechanisms to account for DL population dynamics and especially for swarm movement. For example, Sun *et al*. Sun et al. (2022) used data for elevation, land cover, sand and clay content in soil, soil moisture, NDVI and surface temperature, combined with historical locust ground survey data to produce dynamic forecasts of DL hopper band occurrence. The approach Sun et al. (2022) did not, however, extend to accounting for the migratory behaviour of desert locust swarms that arise from hopper bands, something we address in the integrated modelling framework introduced here. Mamo and Bedane Mamo and Bedane (2021a,b), meanwhile, used deterministic ordinary differential equations to model the effects of early intervention on DL outbreaks and impacts on crop production. The models of Mamo and Bedane Mamo and Bedane (2021a,b) provide valuable mathematical insights into the dynamics of control at given sites, but the models do not account for uncertainty due to weather driven variability nor for swarm movement. Magor *et al*. Magor et al. (2008) also analysed early control intervention in DL campaigns, using historic data and a simple model with a locust multiplication factor dependent on breeding conditions at a given site. In our model, the dynamic conditions of breeding sites are incorporated by recent precipitation events in addition to the intrinsic suitability of soil conditions. We also allow for turbulent wind-assisted dispersal of DL swarms by coupling breeding and emergence of hopper bands with a weather-driven dispersal model, interspersed with variable feeding durations at landing sites.

Migrant pests such as DL could be considered to be metapopulations occupying discrete habitat patches, with colonization and extinction rates defining dynamics of whole population (Cheke and Tratalos, 2007; Ibrahim, 2008). Full parameterisation of a metapopulation model requires data on population sizes within the patches, which are not available for the 2019-2021 upsurge in East Africa. Our approach does not require knowledge of the size of DL population since we model a coherent unit of gregarious locusts at a similar developmental stage. A multi-agent system to represent events of locust plague development and management decisions was designed in (Gay et al., 2017, 2019, 2021). The model was built on a virtual territory represented by a grid of 100 cells, each cell being 10 km by 10 km. Similar to our approach, the main entity comprised coherent groups of locusts. Locust dispersal was simulated by swarms randomly moving to nearby cells, which is in contrast to our approach, where we used meteorological and environmental data to influence swarm movement. Recently, the migration paths of DL during the 2018-2020 upsurge in Africa and Asia were reviewed in conjunction with DL survey data and the distribution of suitable locust habitats (Dong et al., 2023). However, the authors have not provided details on which meteorological products were used to extract wind direction and wind speed, nor how individual migratory paths were constructed.

In our modelling framework we utilised the ability of the NAME atmospheric dispersion model to reproduce stochasticity associated with turbulent wind flow. Essentially this involves modelling an ensemble of possible trajec-tories from a starting location, each of which represents one possible pathway through the turbulent atmosphere. In previous studies on wind-assisted insect migration, the direction of migration was assumed to follow a single air parcel trajectory. One popular choice is to use wind trajectories obtained from the HYSPLIT modelling system Stein et al. (2015). HYSPLIT trajec-tories have been used to study migratory trajectories of many insect species, such as seasonal migration of fall armyworm moths in the USA (Westbrook et al., 2015), and trans-Saharan migration of painted lady birds (Hu et al., 2021). During the 2019-2021 DL upsurge, NOAA created a web-based app for forward or backward simulation of swarm movement based on HYSPLIT’s air trajectories (NOAA). Simulated HYSPLIT forward trajectories were also used to asses DL invasion risk to China by Wang *et al*. Wang et al. (2020). Another alternative approach to derive single air parcel wind trajectories is to use the **u** and **v** values of wind speed components derived from re-analysed meteorological data or from weather forecasting models. The displacement of a swarm of migratory locusts of the species *Schistocerca cancellata* was sim-ulated using the directions and intensities of the wind derived from Weather Research and Forecasting model (Tumelero et al., 2021). The advantage of accounting for the inherent stochasticity of wind-assisted dispersal as in the framework proposed here lies in the ability to compare the likelihood of different trajectories and DL landing sites from an initial site. We note that the HYSPLIT modelling platform can also be used to calculate ensemble pathways.

Our framework follows gregarious locust populations through egg, hopper and adult stages. By separating the breeding process into two parts (breeding suitability map and recent precipitation events that favour egg laying and development) we can take advantage of the high temporal resolution data on temperature, precipitation and soil moisture provided by the UM. We found broad correspondence between our prediction and predictions at broad scales in (Kimathi et al., 2020). In addition to the increased spatial resolution, an advantage of our approach to predict breeding suitability is that elevation, sand and clay content in soil are invariant to climate change, at least for the current time horizon used for climate change impact evaluation.

Feeding capacity was assessed based on land cover type, NDVI trend and vegetation state. Longest stay was assumed for cropland, with higher and highest density of vegetation and with increasing and constant NDVI value, which indicates pre-peak and peak of crop development stage. Further refining of the rules using observational field data on the feeding behaviour of swarms would be beneficial to improve predictability.

Swarms follow seasonal shifts of the Intertropical Convergence Zone (ITCZ), which can take them to areas of rainfall (Rainey, 1951; Draper, 1980; Pedgley, 1981; Homberg, 2015; Homberg et al., 2022). Vallebona *et al*. (Vallebona et al., 2008) showed that the DL upsurge mechanism is linked with a stronger westerly mid-latitude circulation in March followed by a weakened African Easterly Jet and a strengthened moisture advection from April to May. Analysis of the desert locust plague in 1950 indicated a high association between swarm reports and the day after which N-E winds established at the surface for a greater part of each day (Rainey, 1951). We investigated variation in wind patterns by fitting the von Mises distribution to angles representing the direction of NAME wind trajectories. The von Mises distribution is a distribution commonly used for modelling circular data (Mardia, 1975), as well as for statistical modelling of directional wind speeds (Carta et al., 2008). Our analysis confirmed the influence of the ITCZ on local wind trajectories, i.e. southwards orientation in January and northwards orientation in August. Certain areas tended to have a smaller concentration in the direction of wind trajectories, which introduces larger variability in model simulations to predict DL landing sites.

To test the framework, we simulated breeding, development and migration histories for locations and time periods chosen based on the observed field surveillance data. Our simulations showed good correspondence with the observed increase in swarms reaching the northeast of Kenya after 1st December 2020. The modelled spread of swarms is in agreement with previous evidence that swarms may fly up to nine hours and may easily move 200 km or more in a day (Cressman, 1996; Symmons and Cressman, 2001).

Our proposed algorithm is particularly useful to model short-term and long-term desert locust migration patterns with limited reporting data. The projected feeding and migration trajectories could be used to guide surveillance. We used historic wind data from the UK Met Office Unified model to map risk for up to seven days. This limit is set by the current availability of weather *forecast* data for generating future wind trajectories for use in the NAME model. A period of seven days should be sufficient for control operations to be implemented (FAO and WMO, 2016).

In East Africa, the 2019–2021 DL upsurge was characterised by a high degree of spatial and temporal heterogeneity, with some regions experiencing persistently higher DL invasion frequency (Retkute et al., 2021). Our proposed modelling framework can help to explain this heterogeneity, and can help to improve locust monitoring and response efforts. Further work is under way to assess different surveillance methods employed during the 2019–2021 DL upsurge, to perform rigorous parameter estimation of the model, and to investigate how to use the modelling framework to reduce risks from the harmful effects of DL outbreaks by including control measures under resource constraints. This will become more critical as climate change influences the distribution of DL (Meynard et al., 2017) and it is therefore important to be able to predict DL migration in new regions. The intention is to have the modelling framework ready now to be used if another upsurge were to occur.

## Acknowledgments

The author would like to thank Rebekah GK Hinton for constructive criticism of the manuscript.

## Supplementary Information

**Supplementary Table 1:** SI figure 1: Input data.

|  | Type (Static/Dynamic) | Scale | Source | Provider |
| --- | --- | --- | --- | --- |
| Elevation | Static | Global<br>1km x 1km | https://srtm.csi.cgiar.org | The Shuttle Radar Topography Mission (SRTM) digital elevation dataset |
| Sand content | Static | Global<br>250m x 250m | https://files.isric.org/soilgrids/former/2017-03-10/data/ | ISRIC soilgrids dataset |
| Clay content | Static | Global<br>250m x 250m | https://files.isric.org/soilgrids/former/2017-03-10/data/ | ISRIC soilgrids dataset |
| Soil moisture | Dynamic | Ethiopia, Somalia, Eritrea, Kenya, Djibouti<br>10km x 10km<br>Temporal scale: 3h | This project | UK Met Office |
| Land surface temperature | Dynamic | Ethiopia, Somalia, Eritrea, Kenya, Djibouti<br>10km x 10km<br>Temporal scale: 3h | This project | UK Met Office |
| NAME wind trajectories | Dynamic | Ethiopia, Somalia, Eritrea, Kenya, Djibouti<br>Source resolution: 10 km x 10 km<br>Temporal scale: 0.5h | This project | UK Met Office |
| Land cover type | Static | 100m x 100m | https://zenodo.org/record/3939050,<br>doi:10.5281/ZENODO.3939050 | The Copernicus global map of land cover at 100 m resolution (CLC100) |
| NDVI | Dynamic | 250m x 250m<br>16d | https://developers.google.com/earth-engine/datasets/catalog/MODIS_061_MOD13Q1 | NASA LP DAAC at the USGS EROS Center |
| DL reports | Dynamic | Ethiopia, Somalia, Eritrea, Kenya, Djibouti<br>Dates: 2019/01/01-2021/12/31 | https://locust-hub-hqfao.hub.arcgis.com/search?collection=Dataset | FAO Locust Hub |

C.A.G.

**Supplementary Table 2:**
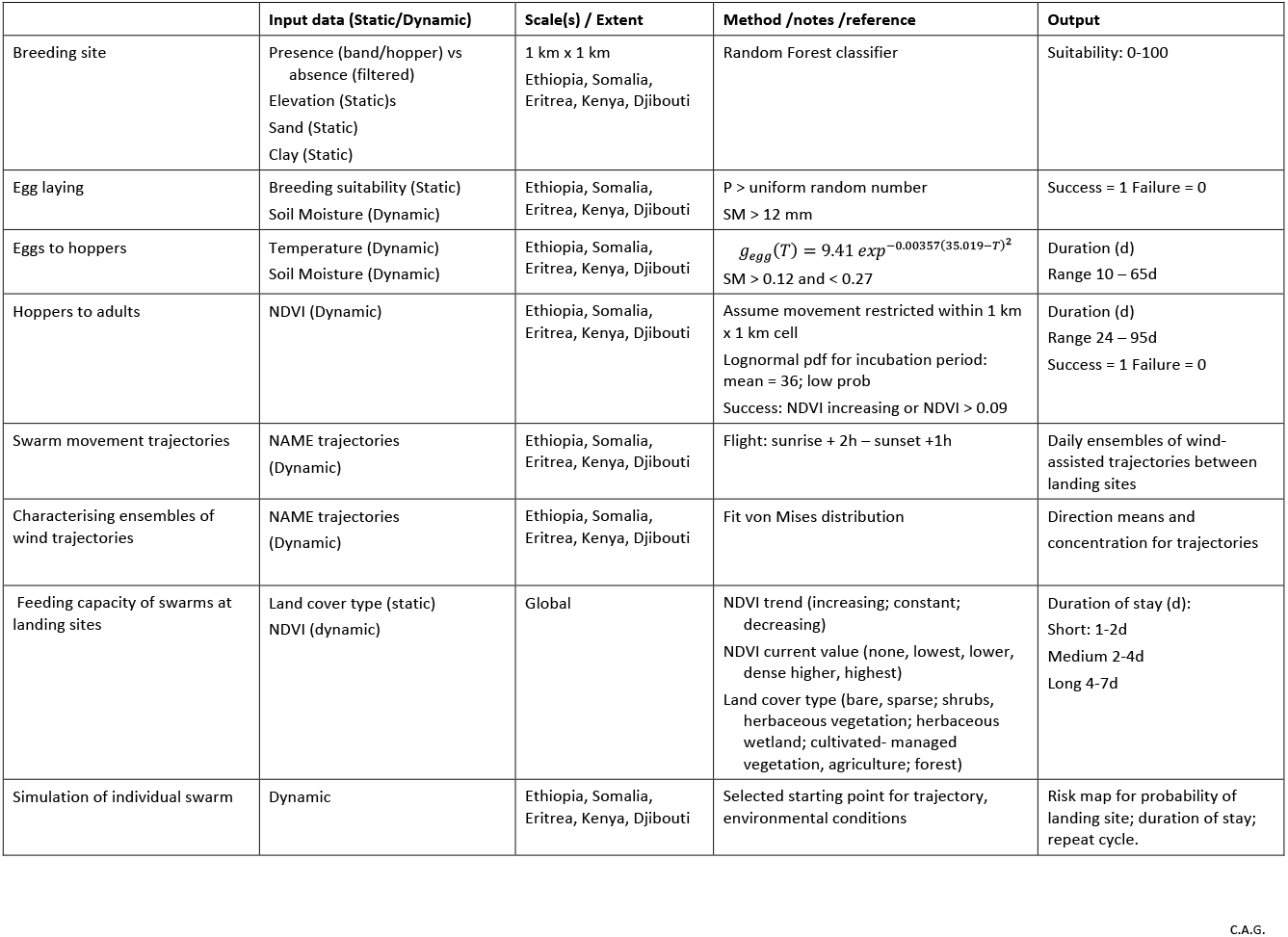
SI figure 2: Summary of the desert locust modelling framework.

